# Peroxisome proliferator-activated receptor gamma (*PPARG*)-mediated myocardial salvage in acute myocardial infarction managed with left ventricular unloading and coronary reperfusion

**DOI:** 10.1101/2025.02.11.637726

**Authors:** Joseph R. Visker, Eleni Tseliou, Christos P. Kyriakopoulos, Rana Hamouche, Michael Yin, Jing Ling, Thirupura S. Shankar, Eugene Kwan, Luis Cedeno-Rosaria, Jesse N. Velasco-Silva, Konstantinos Sideris, Hyoin Kwak, Yanni Hillas, Eleni Yannias, Eleni Maneta, Harini Srinivasan, Laisha Padilla, Georgiy Polishchuck, Sutip Navankasattusas, Anwar Tandar, Gregory S. Ducker, Jared Rutter, TingTing Hong, Robin M. Shaw, Charles Lui, Frederick G. Welt, Stavros G. Drakos

## Abstract

Ischemic heart disease and acute myocardial infarction (AMI) is a leading cause of morbidity and mortality. Improvements have been made in coronary interventions to restore blood flow, but ischemia/reperfusion (I/R) injury significantly impacts clinical outcomes. We previously reported that activation of percutaneous mechanical unloading of the left ventricle (LV) with a transvalvular axial-flow device simultaneously with reperfusion improves myocardial salvage. However, the underlying mechanisms, potential adjuvant pharmacological interventions and the timing of the use of LV unloading as a cardioprotective approach in AMI are not well understood. This study investigated a) the mechanisms associated with improved myocardial salvage, b) a pharmacological intervention, and c) the timing of LV unloading. Following 90 minutes of ischemia, adult swine were subjected to reperfusion alone, simultaneous unloading with reperfusion, upfront unloading with delayed reperfusion, upfront reperfusion with delayed unloading, or reperfusion with concurrent use of esmolol and milrinone. Compared to controls, the simultaneous group had a 47% increase in myocardial salvage following AMI. This was associated with increased expression of neutrophil degranulation, macrophage activation, iNOS signaling, wound healing, and PPAR signaling. From these pathways, *PPARG* (peroxisome proliferator-activated receptor gamma) emerged as a potential cardioprotective gene that was uniquely overexpressed in the simultaneously unloaded and reperfused myocardium. Next, we showed *PPARG* agonism with rosiglitazone reduces mitochondrial oxygen demand in cardiomyocytes and in vivo, improves myocardial salvage following I/R injury in *C57BL6/J* mice. Thiazolidinediones (TZDs), such as rosiglitazone could be investigated as therapies combined with simultaneous LV unloading and coronary interventions to mitigate reperfusion injury.

**GRAPHICAL ABSTRACT:** 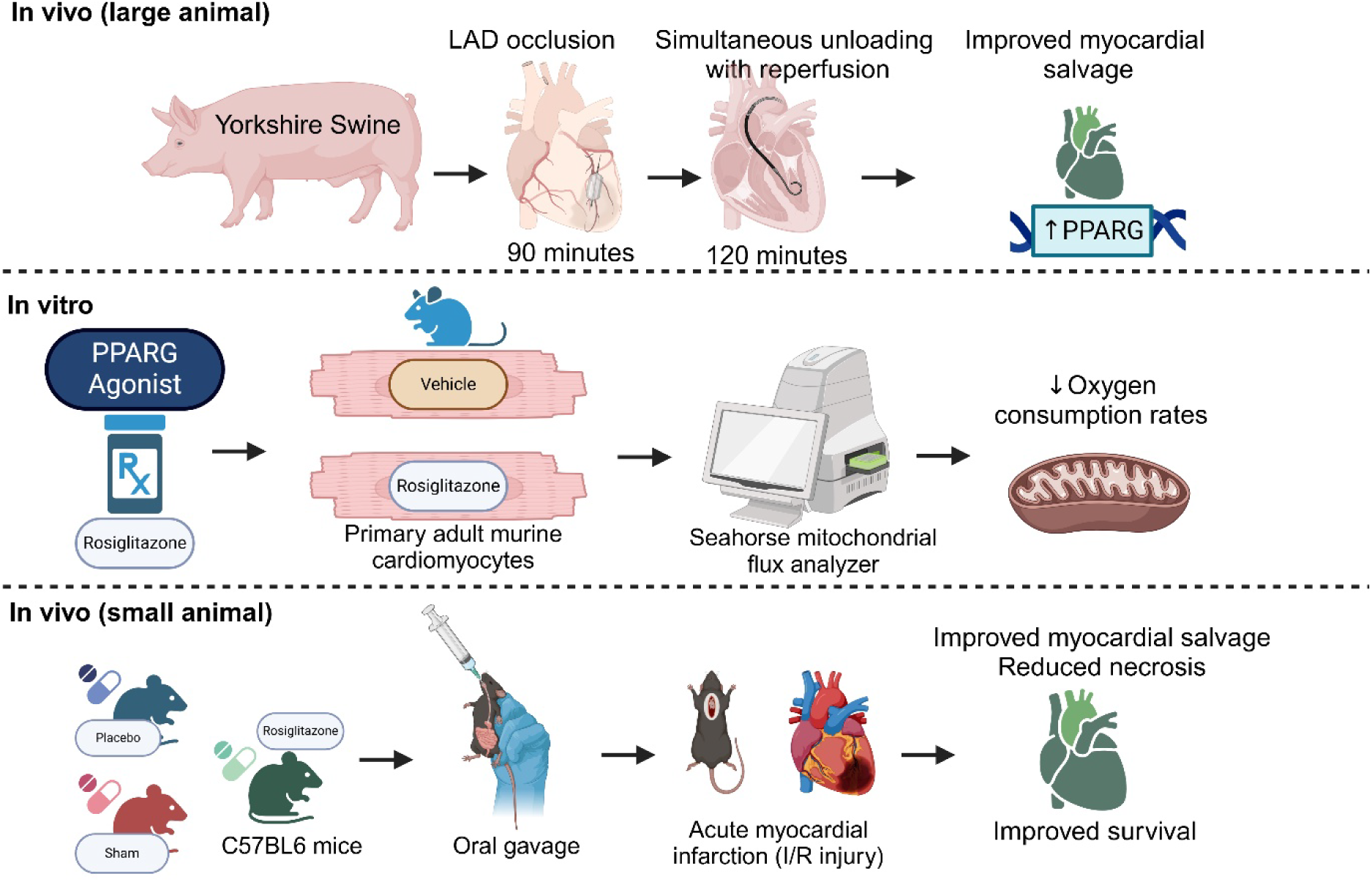

## INTRODUCTION

Acute myocardial infarction (AMI) is a leading cause of mortality worldwide with an estimated 17.9 million deaths per year ^1^. In the U.S. alone, AMI occurs every 2 minutes, which amounts to about 805,000 Americans suffering AMI annually ^2^. AMI is associated with a 15% mortality rate during index hospitalization and up to 25% during the first year, with long-term complications including chronic heart failure, making it a critical public health issue ^3^. The current standard-of-care is primary percutaneous coronary intervention (PPCI), a procedure aiming to restore blood flow to the ischemic myocardium ^1,2,4^. While PPCI indeed restores blood flow and myocardial perfusion which is mostly beneficial, the rapid re-introduction of oxygenated blood causes further injury to the heart, known as “ischemia-reperfusion injury” (I/R) ^5,6^. I/R reduces the clinical benefits of PPCI by sometimes increasing the final infarct size ^7^. The larger the infarct size the higher the likelihood to lead to ischemic cardiomyopathy and chronic heart failure (HF) ^3,8^. Investigating the mechanisms involved in I/R may identify new therapeutic strategies to complement the existing standard-of-care with the goal of improving clinical outcomes for patients suffering AMI ^9^.

We, and others have previously shown that acute percutaneous mechanical unloading of the left ventricle (LV) with a transvalvular axial flow microvascular pump (Impella®, Abiomed) prior to or simultaneous with reperfusion is capable of improving myocardial salvage following I/R ^10–14^. Even though these devices are under active clinical investigation, our mechanistic understanding of how they confer cardioprotection to the injured myocardium remains limited ^14^. These interventions have been suggested to account for improved myocardial salvage through mechanisms aimed at decreasing the hearts oxygen demand, preserving mitochondrial integrity, and inactivating pro-apoptotic pathways leading to cell survival ^12^. Similarly, other groups have proposed investigating pharmacological interventions, such as the combination of esmolol and milrinone, as an alternative cardioprotective strategy to mitigate I/R injury ^15–19^. Esmolol, a short-acting β-blocker, reduces myocardial oxygen demand by decreasing heart rate and contractility while milrinone, a phosphodiesterase inhibitor, promotes vasodilation, potentially improving coronary blood flow. This pharmacological approach may mimic the beneficial effects of LV-unloading with Impella® devices by reducing myocardial stress and preventing cellular apoptosis, thereby improving myocardial salvage following I/R injury.

The main objectives of this study were a) define the molecular signature associated with cardioprotection from I/R, b) determine if pharmacotherapy with esmolol and milrinone could substitute for mechanical unloading, and c) to determine the optimal timing of LV-unloading with reperfusion. To test this, we used a translational approach using a percutaneous, closed-chest swine model of I/R, transcriptomics, in vivo murine cardiac injury, and oxygen consumption rates of primary adult murine cardiomyocytes.

## METHODS

### Sex as a biological variable

Our study examined male and female animals, and similar findings are reported for both sexes.

All animals were treated according to the guidelines provided by the institutional animal care and use committee (IACUC Protocol #22-02006) at the University of Utah. Adult Yorkshire swine were ordered from an approved USDA farm (Premier BioSource), and were given a physical examination by a third-party veterinarian. After arrival to the University of Utah, and another physical examination, all animals were provided with a ten-day quarantine period to acclimate to their new surroundings. If animals were not deemed healthy prior to the experiment they were excluded from the study.

### Anesthesia and Surgical Preparation

On the day of the experiment, animals were premedicated with tiletamine and zolazepam (4.4 mg/kg, intramuscular (IM)), ketamine (2.2 mg/kg, IM), and xylazine (2.2 mg/kg, IM). General anesthesia was induced and maintained with inhaled isoflurane (1%– 5%). Heating pads, blankets, and socks were used to maintain a core body temperature above 99°F. Using real-time ultrasound, bilateral femoral arteries were accessed with 7F sheaths, and the right femoral vein was accessed for infusion of intravenous (IV) medications. A Swan-Ganz catheter was placed in the pulmonary artery under fluoroscopy via a 9F right internal jugular (IJ) venous sheath.

### Baseline Measurements and Drug Administration

After obtaining baseline hemodynamic measurements, 300 mg of amiodarone was administered IV over 30 minutes, followed by an IV infusion at 1 mg/min. Lidocaine was administered IV at 1 mg/kg over 1-2 minutes, followed by a second dose of 0.5 mg/kg IV over 1-2 minutes 20 minutes later, and maintained at 1 mg/min continuous infusion throughout the experiment. Unfractionated IV heparin was used to maintain an activated clotting time of 250-300 seconds.

### Ischemia and Reperfusion (swine)

Under fluoroscopy, a 6F left Amplatz-1 coronary guide catheter (Medtronic) was used to engage the left coronary artery via the right femoral arterial sheath. A 0.014-inch coronary wire was advanced into the left anterior descending artery (LAD), visualized with contrast dye (Visipaque). A 3.5 mm over-the-wire semi-compliant coronary balloon was advanced to the mid-LAD, just distal to the first diagonal branch and inflated to nominal pressure to occlude coronary blood flow. A coronary angiogram was then performed to ensure complete cessation of blood flow distal to the balloon.

At the end of the ischemic period (90 minutes), the balloon was deflated and retrieved into the guiding catheter. Several coronary angiograms were performed to ensure adequate coronary blood flow was restored to the entire LAD during reperfusion. Cardiac performance was assessed by invasive hemodynamics at baseline, halfway through ischemia, and halfway through reperfusion using the percutaneous Swan-Ganz catheter. Additionally, blood samples were collected at baseline, halfway through ischemia, and halfway through reperfusion.

### Left Ventricular Unloading

The Impella CP with smart assist device (Abiomed) was inserted via a standard catheterization procedure through the femoral artery, into the ascending aorta, across the valve, and into the LV. The device pulls blood from the LV through an inlet area near the tip and expels blood from the catheter into the ascending aorta. To determine the optimal timing of unloading, the Impella device was activated before, after, and simultaneous with reperfusion.

### Hemodynamic Monitoring

To assess the hemodynamic effectiveness of LV unloading, 9 Fr vascular sheaths were used to access the right carotid artery and jugular vein. A Swan-Ganz catheter was advanced through the jugular vein to measure invasive hemodynamics including right- and left-sided filling pressure, cardiac output and pulmonary and systemic vascular resistance.

### Thoracotomy and Tissue Staining (swine)

At the end of the experiment, a thoracotomy was performed, and another 6F JR catheter was introduced through the left femoral artery sheath to engage the right coronary ostium. The coronary balloon was then inflated at the same location (mid-LAD) and 150 ml of 4% Evans Blue (Sigma-Aldrich) was injected into both coronaries to stain the whole myocardium except for the area at risk. Within 10 seconds of the injections, fatal ventricular fibrillation was induced, and the heart was harvested. Five parallel transverse sections of 1 cm thickness were made from the apex to the base of the left heart. The samples were stained with 2,3,5-triphenyltetrazolium chloride (TTC) to demarcate salvageable myocardium. Both sides of each sectioned sample were photographed to quantify the injury to the heart using ImageJ software and quantified by independent researchers who were blinded to the treatment groups using ImageJ (NIH software).

### Ischemia and Reperfusion (mouse)

We used a previously published model of I/R injury with 12-14 weeks old *C57BL/6J* male and female mice weighing between 20-30 grams ^20^. Briefly, mice were anesthetized with 2-3% isoflurane and placed in the supine position on a circulating warming pad (38°C). Next, mice were orally intubated with a 22 GIV catheter, and ventilated using a small rodent respirator set at 100 breaths/minute with a tidal volume adjusted to 15-20 mmHg (Harvard Instrument). A thoracotomy was performed to visualize the anterior surface of the heart. The pericardium was removed using forceps and scissors. Next, the LAD was ensnared with a 6-0 silk suture. A double knot was then made with the suture, leaving a 2-3 mm diameter loop through, then a 2-3 mm long piece of PE-10 tygon tubing was placed into the knot. The loop around the artery and tubing was tightened for 30 minutes to ensure ischemic injury to the myocardium as determined by the paler color of the anterior wall. After the ischemic period, the knot was untied to allow reperfusion for 2 hours. After the I/R injury, 3% Evans blue was injected into the left atrium, hearts were excised, and sliced into 1-mm-thick transverse sections. Next, the slices were incubated (37°C) in 1% TTC to stain the non-infarcted myocardium. Then slices were placed in 10% formalin, imaged, and quantified by independent researchers who were blinded to the treatment groups using ImageJ (NIH software).

### RNA isolation procedure

RNA yield and purity (A260/A280 > 2.0) were assessed using a NanoDrop™ Lite Spectrophotometer (Thermo Scientific™). The RNA-sequencing library was prepared with the Illumina TruSeq Stranded RNA kit with Ribo-Zero Gold to remove ribosomal RNA and sequenced on an Illumina HiSeq 2500 with 50 bp single-end reads. RNA sequence reads were aligned to the swine genome (*sus scrofa domesticus)* version Sscrofa11.1 using Novoalign (version 2.8.1) against an index containing genome sequences plus all splice junctions (known and theoretical, 46 bp radius) generated with USeq MakeTranscriptome (version 8.8.1) and Ensembl gene annotation (release 84), allowing up to 50 alignments per read. Raw alignments were converted to genomic coordinates using USeq 9. SamTranscriptomeParser (version 8.8.8), permitting a maximum of 1 alignment per read. RNA-Seq quality was evaluated using Picard CollectRnaSeqMetrics (version 1.137). Gene counts were collected with SubRead featureCounts (version 1.5.1) and Ensembl Sscrofa11.1 annotation (release 87) in a stranded fashion, assigning reads to genes with the largest overlap. Low and non-expressed transcripts were filtered out when the maximum observed count across all samples was ≤ 10 counts. Differentially expressed transcripts were identified using DESeq2 (version 1.16.1).

### Transcriptomics of salvaged myocardium following I/R with LV-unloading

As previously published, RNA-seq analysis was performed in collaboration with the High-Throughput Genomics and Bioinformatics Analysis Shared Resource at the Huntsman Cancer Institute, University of Utah ^21,22^. For the analysis, the swine GRCm38 FASTA and GTF files were obtained from Ensembl release 96. The reference database was built using STAR version 2.7.0f, with splice junctions optimized for 50 base-pair reads. Optical duplicates were eliminated from paired-end FASTQ files utilizing BBMap’s Clumpify utility (v38.34), and adaptors were trimmed from reads using cutadapt 1.16. The trimmed reads were then aligned to the reference database in two-pass mode using STAR, resulting in a BAM file sorted by coordinates. Mapped reads were assigned to genes annotated in the GTF file with featureCounts version 1.6.3. Outputs from cutadapt, FastQC, Picard CollectRnaSeqMetrics, STAR, and featureCounts were compiled and analyzed for sample outliers using MultiQC.

To identify differentially expressed genes, DESeq2 version 1.24.0 was used, focusing on genes with at least 95 read counts across all samples for experimental group. Genes were filtered based on the criteria of FDR < 0.01 and absolute log2 fold change > 0.585 (fold change > 1.516). Enriched GO terms for up- and down-regulated gene sets in each group were identified using Gprofiler, applying a Benjamini-Hochberg FDR p-value correction threshold of < 0.01. For heatmap generation, RNA-seq read counts were RPKM normalized. Heatmaps were created by gene-standardizing the RPKM values (mean = 0, standard deviation = 1) and plotted using XPRESSplot v0.2.2, Matplotlib, and Seaborn.

### Quantification of molecular targets associated with cardioprotection from I/R

As described above, total RNA was isolated from swine heart tissue that was biopsied following the I/R procedure. We used the RNeasy Mini kits (Qiagen) following the manufacturer’s instructions. Complementary DNA (cDNA) synthesis was carried out using a cDNA Reverse Transcriptase Kit (New England Biolabs). TaqMan-based real-time quantitative polymerase chain reactions (qRT-PCR) were then performed on a QuantStudio 7 Pro Real-Time PCR System (ThermoFisher). The housekeeping gene vinculin served as an internal control for cDNA quantification and normalization of the amplified gene products.

### Isolation of primary adult cardiomyocytes

As previously published, mice were sedated, hearts excised and affixed to an aortic cannula^20,21^. Hearts were then perfused with buffers held at 37°C, pH 7.3, and a flow rate of 2 mL/min). Enzymatic digestion was performed for 15 minutes using protease (0.1 mg/mL) and collagenase (1 mg/mL). Next, the heart was perfused with a “stop buffer” (20% serum and 0.2 mM CaCl_2_). The heart was then minced and filtered through a nylon mesh (125 μm). Cardiomyocytes were then sedimented in increasing amounts of calcium chloride (125 μM, 250 μM, and 500 μM) to yield calcium tolerant cells. Following this, the cells were plated onto poly-L-lysine coated dishes in human plasma-like media (HPLM). Cells that were used in the study were rod shaped, with defined striations, and did not spontaneously contract.

### Seahorse mitochondrial substrate oxidation stress test

The Seahorse Mitochondrial Substrate Oxidation Stress Test was performed on a Seahorse XF Pro Analyzer. The day of the experiment, adult cardiomyocytes were plated in 8-wells of a 96-well seahorse plate in Seahorse XF RPMI media (pH 7.4) (without glucose, pyruvate, glutamine or lactate), and pretreated for 1 h with DMSO (vehicle) or 10 μM Rosiglitazone. The Seahorse Mitochondrial Substrate Oxidation Stress Test was performed by acutely injecting 2 mM lactate or 2 mM pyruvate in cells treated with DMSO or Rosiglitazone, with a standard protocol consisting of five measurement cycles for baseline (3 min mixing, 0 min waiting and 3 min measuring) and ten measurement cycles for acute substrate injection (3 min mixing, 0 min waiting and 3 min measuring). After the assay, data was analyzed in the Seahorse WAVE software through the XF Sub Ox Stress Test Report.

### Statistical Analysis

For our statistical analysis, we used GraphPad Prism software, version 10.0.0. The Shapiro-Wilk and D’Agostino-Pearson omnibus tests were employed to check the normality of the data. Significant outliers were identified using ROUT testing and were excluded before further analysis. Data are presented as either mean ± standard deviation (SD) or ± standard error of the mean (SEM). All comparative analyses were conducted using 2-tailed tests. For comparisons between two groups, we used the unpaired Student’s t-test and the two-tailed Mann-Whitney test. Multiple t-tests and 1- or 2-way ANOVAs were utilized for multiple group comparisons. A significance level of p < 0.05 was established *a priori*. If needed, Tukey’s honest significant difference post-hoc test was applied for multiple comparisons.

## RESULTS

### Left ventricular unloading with reperfusion optimizes hemodynamics, improves myocardial salvage, and mitigates injury

We used age matched, adult male and female yorkshire swine to investigate the role of LV-unloading in the myocardial response to I/R (Figure 1A). Body mass was comparable for all groups, however UPFRONT (85.07±12.08 kg, n=3) weighed significantly more compared to CONTROL (61.80±1.90 kg, n=6), (Figure 1B, p=0.02). We subjected these animals to in vivo myocardial I/R injury, recorded hemodynamics, then assessed the extent of myocardial damage. Overall, our hemodynamic data show that LV-unloading sustained heart rate and arterial blood pressure within normal range, and reduced pulmonary capillary wedge pressure following acute I/R injury (Supplemental Figure 1). Using the double-staining histological technique of Evans blue and TTC, we controlled for infarct size and found that the area at risk (AAR) was indistinguishable between all groups, indicating a similar initial ischemic insult (Figure 1C). However, when normalized to the AAR, the myocardial salvage was significantly increased (54.4%, p=0.02) only in the pigs that experienced LV-unloading simultaneous with reperfusion (SIMULT: 71.10±3.08%) when compared to the reperfusion only pigs (CONTROL: 46.06±4.90%), (Figure 1D-F). The ESM+MIL group showed a trend (60.69±6.673, p=0.08) towards improved myocardial salvage when compared to controls, however this finding was not statistically significant. All other groups myocardial salvage was not significantly different when compared to controls (p>0.05).

**Figure 1.**
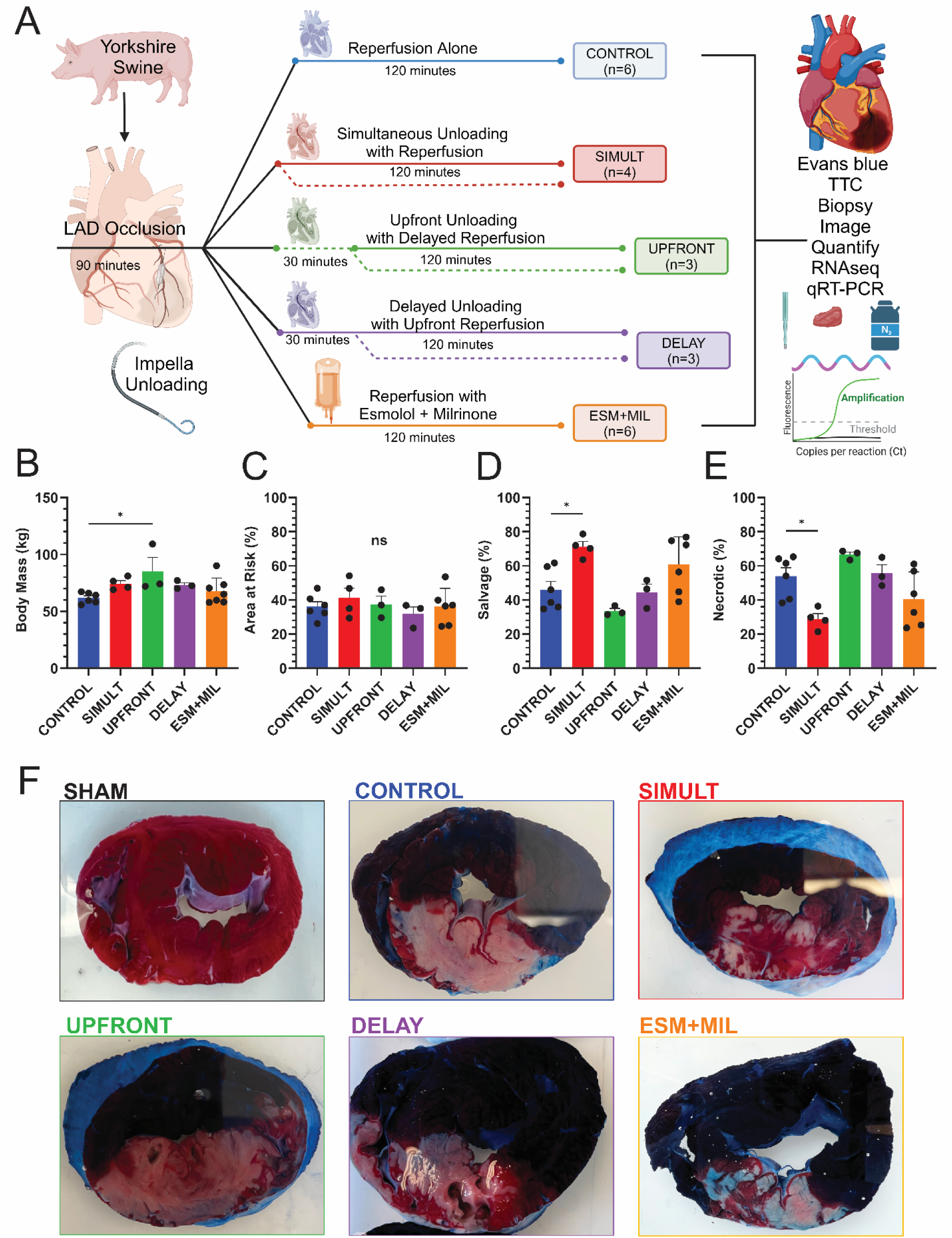
Simultaneous left ventricular unloading and reperfusion myocardial salvage. **A)** Large animal swine model of myocardial I/R injury using percutaneous balloon occlusion of the left anterior descending artery. **B)** Body mass showing UPFRONT (85.07±12.08 kg, n=3) weighed significantly more compared to CONTROL (61.80±1.90 kg, n=6), (p=0.02). All other groups were comparable in mass (SIMULT: 73.88±2.15 kg, n=4; DELAY: 73.20±2.043 kg, n=3; ESM+MIL: 67.80±4.35 kg, n=6). **C)** Area at risk (AAR) was similar between all groups (CONTROL: 36.07±2.92%, SIMULT: 41.32±5.614%, UPFRONT: 37.53±4.72%, DELAY: 31.76±4.15%, ESM+MIL: 36.08±4.38%, p>0.05). **D-E)** Myocardial salvage standardized to AAR was increased in SIMULT (71.10±3.08%) when compared to CONTROL (46.06±4.90%). In all other groups myocardial salvage was non-significant (p>0.05) when compared to controls (UPFRONT: 33.49±1.579%, DELAY: 44.35±4.87%, ESM+MIL: 63.87±7.19%). **F)** Representative images showing myocardial salvage and necrosis in transverse tissue sections stained with Evans blue and TTC. A porcine heart from an animal that was medicated, anesthetized, and intubated but did not experience any I/R injury was also stained (SHAM, n=1). This animal died due to complications with the intubation process.

### Acute ischemia-reperfusion injury causes changes in gene expression

Encouraged by our functional results, we sought tounderstand the underlying mechanisms driving the differential response to I/R injury in the pigs subjected to unloading by performing quantitative transcriptomics on hearts. All groups were subjected to I/R and following two hours of reperfusion, we biopsied the salvaged myocardium before RNA isolation and sequencing (Figure 2A). To detect differentially expressed genes, we utilized a 5% false discovery rate with DeSeq2 software (version 1.24.0), and genes were screened with an adjusted p-value <0.05, and absolute log_2_ fold change > 0.585.

**Figure 2.**
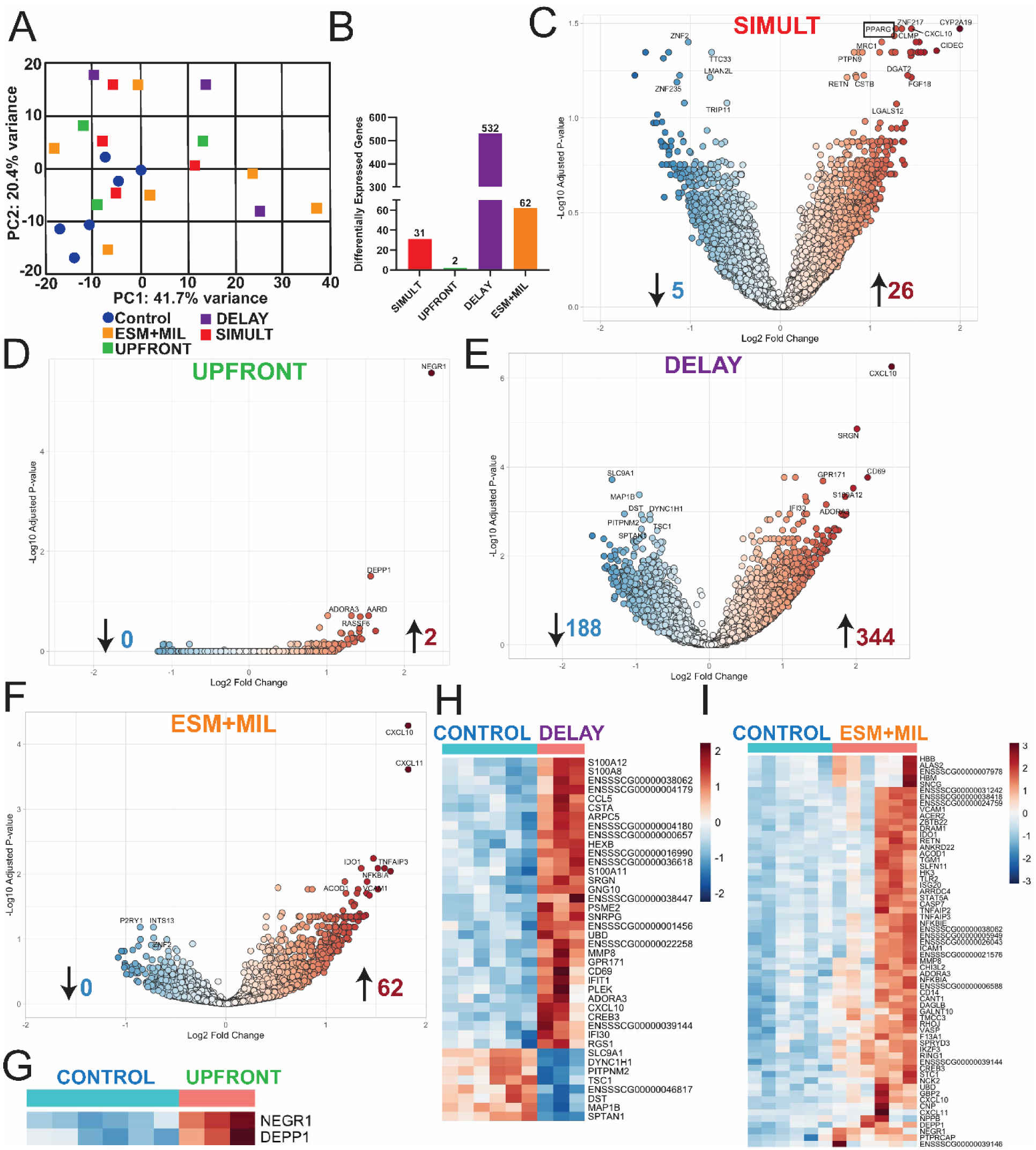
Changes in gene expression associated with acute ischemia-reperfusion injury. **A)** Principle component analysis showing the variability between groups. **B)** Number of differentially expressed genes when compared to the CONTROL group. **C-F)** Volcano plots of differentially expressed genes. **G-I)** Heatmaps of overall transcriptomic signature amongst the UPFRONT, DELAY, and ESM+MIL groups, however these groups did not show an increased myocardial salvage following I/R injury.

In response to I/R injury, large changes in gene expression were identified (Figure 2B). There were 532, 62, 31, and 2 genes differentially expressed with a significant change (p<0.05) in the DELAY, ESM+MIL, SIMULT, and UPFRONT groups, respectively, when compared to CONTROL (Figure 2C-I).

### Simultaneous left ventricular unloading with reperfusion upregulates cardioprotective pathways upon ischemia-reperfusion

Since the SIMULT group showed the most myocardial salvage following I/R injury in our histological results, we chose to focus on the genetic signature in this group leading to cardioprotection (Figure 3A). First, we performed ingenuity pathway analysis (IPA) as a bioinformatics approach to classify enriched groups of genes among those that were differentially expressed in SIMULT compared to CONTROL (Figure 3B). We note that the top canonical pathways that are upregulated in the SIMULT group included neutrophil degranulation (p=3.16E^-21^), macrophage activation (p=3.16E^-14^), phagosome formation (p=2.00E^-10^), and interleukin signaling (p=3.16E^-9^). The top upregulated bio-functions include genes associated with cell function and maintenance (p=5.01E^-31^), cellular movement (p=1.26E^-30^), cell to cell signaling (p=3.98E^-23^), and cell survival (p=3.16E^-18^). We subsequently cross-referenced the 31 significantly identified genes (p<0.05) from the SIMULT group with those identified in the other unloaded hearts. This analysis revealed a subset of six genes uniquely specific to the SIMULT group (Table 1 and Figure 3C). We then performed a STRING protein network analysis and identified that *TRARG1, PLPP3*, and *TNFAIP2* are associated with *PPARG* (Figure 3D). Based upon this information, we then confirmed using qRT-PCR that *PPARG* is significantly upregulated in the SIMULT group when compared to CONTROL and the ESM+MIL group, and no other groups showed this genetic signature (p>0.05) (Figure 3E).

**Figure 3.**
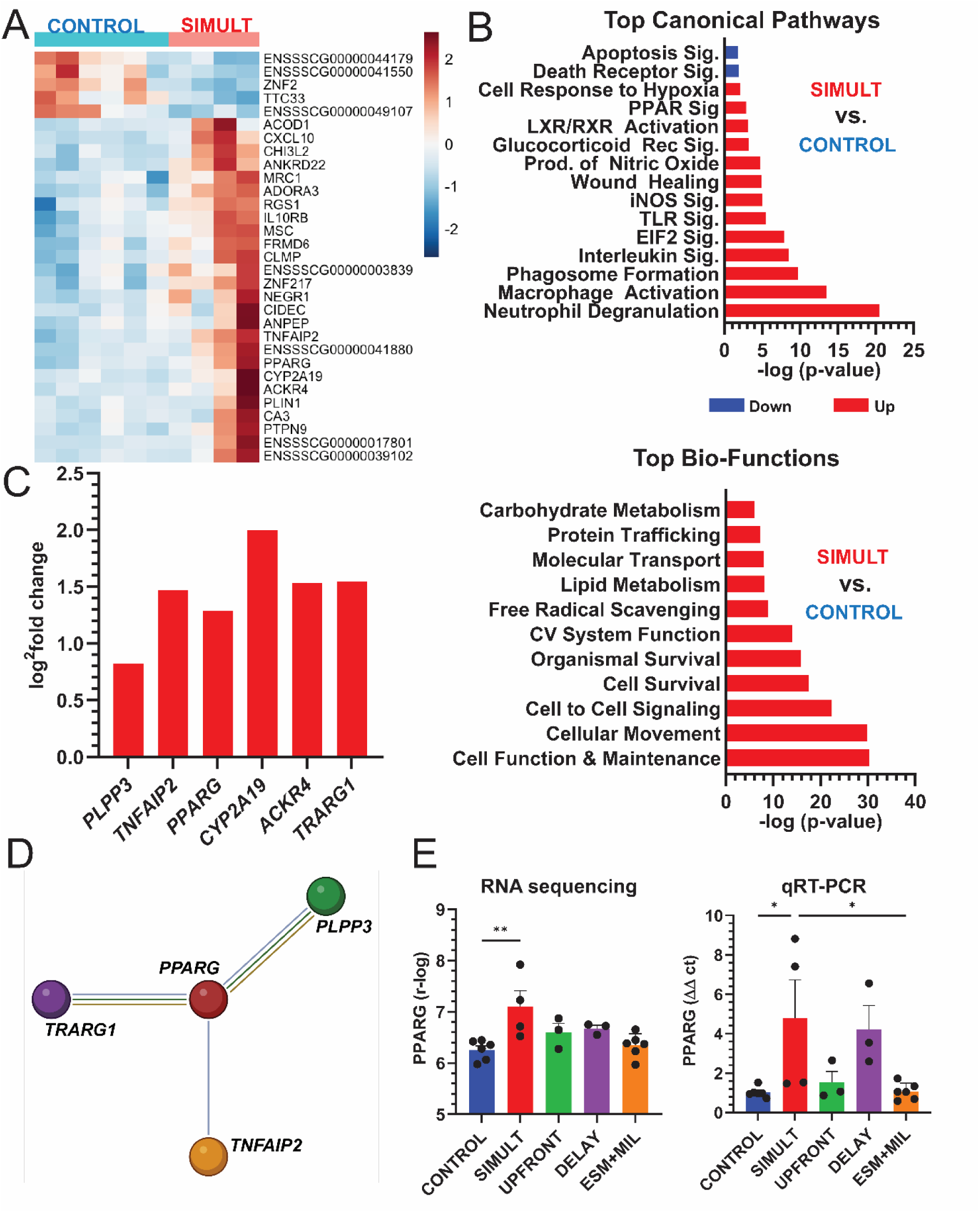
Simultaneous left ventricular unloading and reperfusion upregulates several cardioprotective pathways, including *PPARG* upon ischemia-reperfusion. **A)** Heatmap and names of significantly identified genes in the SIMULT myocardium compared to CONTROL. **B)** Top canonical pathways and bio-functions identified through ingenuity pathway analysis. **C)** Significantly identified genes that are exclusive to the SIMULT group after cross-referencing all genes identified in the unloaded hearts. **D)** STRING protein-protein pathway analysis of the exclusive SIMULT genes showing the nodes focus on *PPARG*. **E)** Standardized transcriptomics r-log values and qRT-PCR data showing significant upregulation of *PPARG* in the salvaged myocardium of SIMULT pigs.

**Table 1.** Genes exclusively expressed in simultaneously unloaded and reperfused pigs.

| Gene name | Symbol | Log<sup>2</sup> fold change | Gene type | Function | Also known as |
| --- | --- | --- | --- | --- | --- |
| Phospholipid Phosphatase 3 | *PLPP3* | 0.82 | Protein coding | Lipid synthesis, and signal transduction | *LPP3*, *VCIP*, *Dri42*, *PAP2B*, *PPAP2B* |
| TNF alpha induced protein 2 | *TNFAIP2* | 1.46 | Protein coding | Mediator of inflammation and angiogenesis | *B94*, *EXOC3L3* |
| Peroxisome proliferator activated receptor gamma | *PPARG* | 1.28 | Protein coding | Transcription factor for β-oxidation and glucose homeostasis | *GLM1*, *CIMT1*, *NR1C3* |
| Cytochrome P450 family 2 subfamily A member 19 | *CYP2A19* | 1.99 | Protein coding | Xenometabolism | *CYP2A*, *CYP2A6* |
| Atypical chemokine receptor 4 | *ACKR4* | 1.52 | Protein coding | G protein-coupled receptor activity and chemokine receptor activity | *PPR1*, *CCBP2*, *CCR11*, *VSHK1* |
| Trafficking regulator of GLUT4 | *TRARG1* | 1.54 | Protein coding | Glucose import and protein transport | *BEC-1*, *DSPB1*, *LOST1*, *TUSC5*, *IFITMD3* |

### PPARG improves myocardial salvage in a murine model of ischemia-reperfusion injury

After identifying *PPARG* as a potential modulator of cardioprotection from I/R injury with activation in our large animal model, we then utilized an in vitro approach to understand whether *PPARG* agonism by rosiglitazone (ROSI) may influence cardiac metabolism. ROSI in the setting of I/R injury may reduce myocardial oxygen demand ^23,24^ and apoptosis ^25^, while supporting cardiac energetics by enhancing fatty acid oxidation and glucose utilization ^26^, thereby mitigating tissue damage. Briefly, we treated culture murine primary adult cardiomyocytes with either vehicle (DMSO) or ROSI (10 μM) and measured mitochondrial oxygen consumption rates (OCR) when provided with either pyruvate or lactate. For both substrates, ROSI-treated cells (pyruvate: 22,362±973.6 auc, lactate: 26,205±811.1 auc) showed a trend toward reduced OCR when compared to vehicle-treated cells (pyruvate: 25,282±1237 auc, lactate: 22,362±973.6 auc) (Figure 4A and B).

**Figure 4.**
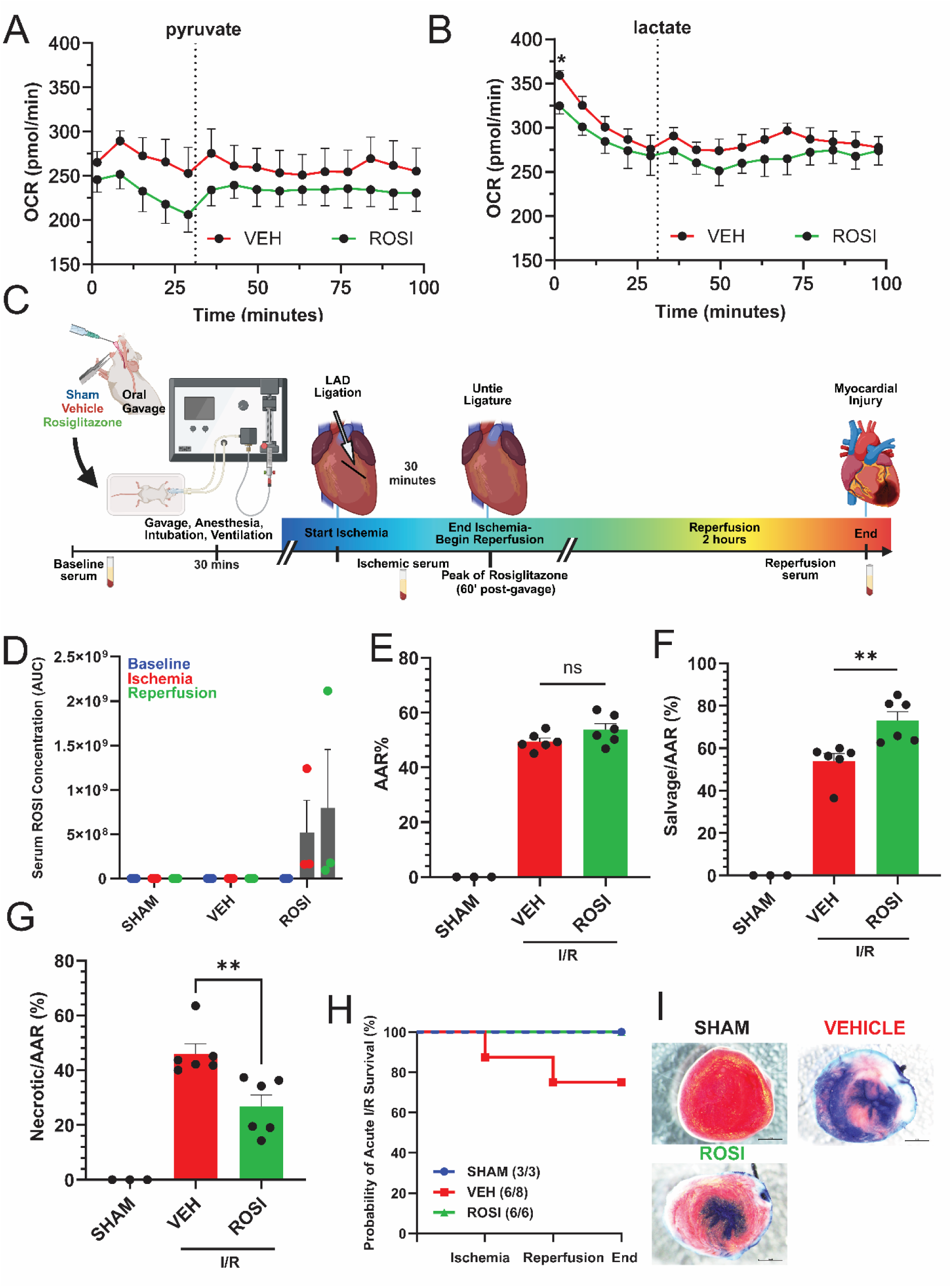
*PPARG* activation with rosiglitazone improves myocardial salvage following ischemia-reperfusion injury in mice. **A and B)** Oxygen consumption rates in primary adult cardiomyocytes introduced to vehicle (DMSO) or rosiglitazone (10 μM) respiring in the presence of pyruvate or lactate. **C)** *C57BL/6* mice were gavaged with either a vehicle (VEH: DMSO, n=6) or rosiglitazone (ROSI, 3 mg/kg, n=6) and exposed to in vivo acute I/R injury. SHAM (n=3) were anesthetized, intubated, and ventilated but but did not experience any cardiac injury. **D)** Serum concentrations of rosiglitazone was detected in the ROSI group following oral gavage at the end of ischemia (1 hour post gavage), and the end of reperfusion (3 hours post gavage). **E)** AAR% was non-significant between vehicle and ROSI groups (p>0.05). **F and G)** Myocardial salvage and necrosis standardized to AAR% is significantly changed in the ROSI treated mice when compared to vehicle. **H)** Vehicle treated mice had reduced survival rates when compared to ROSI treated mice and SHAM controls, but is non-significant (p>0.05). **I)** Representative images showing increased myocardial salvage in the ROSI treated mice compared to vehicle, while SHAM mice show no cardiac injury.

Next, we hypothesized that *PPARG* agonism induced by rosiglitazone would increase myocardial salvage following acute I/R injury. To test this, we subjected *C57BL/6* mice to a single oral gavage of either vehicle control (DMSO) or rosiglitazone (ROSI: 3 mg/kg) and then performed an in vivo acute myocardial I/R procedure. As previously published, the LAD was ligated for 30 minutes and then the ligature was released allowing the myocardium to reperfuse for two hours ^20^. Following the acute I/R procedure, the myocardium was stained with Evans blue and TTC to visualize and quantify the myocardial injury (Figure 4C). We also included a sham-control group (n=3). Mice subjected to the sham operation were anesthetized and intubated but experienced no myocardial I/R injury. We verified via LC-MS that ROSI was present in the serum post-oral gavage at the end of ischemia and reperfusion (Figure 4D). Similar to our swine model, we controlled for infarct size and found that the AAR was comparable between vehicle control (49.46±1.27%) and ROSI (53.80±2.16%) treated mice (p=0.19), (Figure 4E). Yet, when standardized to the AAR, myocardial salvage was increased by 35% in the mice given a single dose of ROSI (73.12±4.13%) when compared to vehicle treated mice (53.94±3.56%), (p=0.006), (Figure 4F and G). Additionally, we dobserved a reduced probability of survival in the vehicle treated mice compared to the ROSI and SHAM mice, however this finding did not reach statistical significance (p>0.05), (Figure 4H and I).

## DISCUSSION

Despite improvements in implementing prompt and timely revascularization, AMI and coronary artery disease remain the largest cause of mortality and inciting event for developing chronic heart failure worldwide ^27–30^. Currently, there is an unmet clinical need to develop novel pharmacological and/or mechanical therapies to mitigate reperfusion injury after coronary intervention for patients suffering AMI. The use of LV unloading with an endovascular miniature heart pump (for example the Impella device or other similar emerging technologies) has been proposed as a potential cardioprotective strategy to improve survival from reperfusion injury. It has been suggested that LV unloading during PPCI after AMI can reduce myocardial wall stress ^31,32^, attenuate oxidative stress ^33^, improve coronary perfusion ^11^, and protect mitochondrial function ^34^. Together these mechanisms work synergistically to protect the myocardium from the damaging effects of reperfusion injury. However the optimal timing to deploy LV unloading is under investigation ^10–14,35–37^.

In this study, our group reproduced our previously reported findings ^10^ and confirmed that LV unloading coupled simultaneously with reperfusion yields the largest amount of myocardial salvage. Based upon the reproducibility of our findings, we hypothesized that we could determine the cardioprotective biology associated with simultaneous unloading with reperfusion. Additionally, we investigated the use of esmolol and milrinone as a pharmacological alternative to mechanical LV unloading. The use of esmolol and milrinone has been shown to reduce I/R injury in rodents ^15,17–19^, presumably by reducing myocardial oxygen demand and improving coronary perfusion without the risks associated with deploying mechanical devices such as valvular injury ^38^, thromboembolism or other vascular complications ^39^.

### Simultaneous unloading with reperfusion decreases ischemia-reperfusion injury

We subjected adult Yorkshire pigs to acute I/R injury with 90 minutes of ischemia followed by 120 minutes of reperfusion. To determine the ideal timing of LV unloading we randomized our pigs to five groups. 1) *CONTROL*: reperfusion with no Impella unloading (n=6), 2) *SIMULT*: Simultaneous unloading with reperfusion (n=4), 3) *UPFRONT*: Upfront unloading with delayed reperfusion (n=3), 4) *DELAY*: Upfront reperfusion with delayed unloading (n=3), and 5) *ESM+MIL*: Reperfusion with use of esmolol and milrinone (n=6). SIMULT animals had >50% increase in myocardial salvage when compared to the control group of PCI reperfusion only, which is currently the standard clinical care for AMI patients (Figure 1). Of note, we observed the ESM+MIL animals showing a trend (p=0.08) towards improved myocardial salvage following I/R injury when compared to controls, however this did not reach statistical significance. The UPFRONT animals, which underwent 30 minutes of unloading prior to reperfusion showed the least amount of myocardial salvage. Following transcriptomics of the salvaged myocardium, we observed an upregulation of several cardioprotective pathways and transcripts after I/R injury (Figure 2), except in the UPFRONT myocardium which only revealed two significant genes (*NEGR* and *DEPP1*). Using the improved myocardial salvage as a condition for further downstream analysis, we investigated the unique genetic signature of the SIMULT group. While our analysis does not definitively establish these genes as the most critical for understanding the enhanced salvage in the SIMULT group, their unique identification suggests a strong likelihood of relevance. The top canonical pathways identified were neutrophil degranulation, macrophage activation, and phagosome formation (Figure 3). Together, these pathways may coordinate a controlled inflammatory response that clears damaged cells and debris while limiting excessive tissue damage^40–47^. Neutrophils can release protective enzymes and cytokines that modulate inflammation^48,49^, while activated macrophages and phagosomes help remove apoptotic cells and promote tissue repair^50,51^, collectively reducing the myocardial injury and enhancing recovery.

The improved myocardial salvage was also associated with an increased expression of peroxisome proliferator-activated receptor gamma (*PPARG*). *PPARG* is crucial for maintaining the metabolic flexibility and functional integrity of the myocardium protecting it from metabolic dysregulation, inflammation, and oxidative stress which are key factors in reperfusion injury ^24–26,52–56^. It is a transcription factor that, upon activation, forms a heterodimer with another nuclear receptor, *RXR* (retinoid X receptor). The *PPARG-RXR* complex modulates the expression of genes involved in glucose and lipid metabolism. Thus, *PPARG* is uniquely situated as it regulates the expression of genes involved in fatty acid uptake, transport, and oxidation while also promoting insulin sensitivity and glucose uptake into cardiomyocytes. Additionally, *PPARG* has been shown to reduce oxidative stress due to reactive oxygen species (ROS) production ^26,57–59^. The upregulation of *PPARG* in the salvaged myocardium of the SIMULT hearts suggests a cardioprotective metabolic program capable of sustaining fatty acid oxidation and glucose metabolism during I/R injury, which was not observed in our other experimental groups.

### PPARG activation can reduce ischemia-reperfusion injury

Thiazolidinediones (TZDs), like rosiglitazone (ROSI) are a class of drugs used to improve insulin sensitivity in patients with type II diabetes ^60–62^. Agonists of *PPARG* have been used previously in the setting of I/R injury with varying results ^53–55,57–59,63^. However, the majority of evidence suggests they are cardioprotective to the myocardium under stressful conditions, such as hypertension, arrhythmias, and ischemia.

First, to understand how *PPARG* activation influenced mitochondrial metabolism, we treated isolated primary adult cardiomyocytes (ACMs) with either a vehicle (DMSO) or ROSI (10 μM) and then exposed them to different energetic fuels, pyruvate or lactate. We noticed that the ROSI treated ACMs, in general, had lower oxygen consumption rates (OCR) when compared to the vehicle-treated cells, suggesting a reduced mitochondrial oxygen demand. From this, we hypothesized that ROSI treatment would be sufficient to drive myocardial salvage, and we tested this hypothesis using an in vivo acute I/R mouse model. We showed that a single dose of ROSI administered via oral gavage (3 mg/kg) before I/R resulted in a 35% increase in myocardial salvage following injury when compared to vehicle-treated mice.

These results show that *PPARG* activation can be cardioprotective following I/R injury. Our data suggest that activating *PPARG* has the potential to reduce mitochondrial oxygen demand and alleviate acute I/R injury. We hypothesize this may in part be due to the ability for *PPARG* to sustain the metabolic choice between glucose and fatty acid metabolism during stressful conditions. This cardioprotective mechanism may bolster the myocardium from oxidative stress and sustain the transition towards metabolic flexibility after I/R injury, however more research is needed to support this hypothesis.

### Conclusions and Perspectives

LV unloading with an endovascular heart pump during AMI is a reproducible and feasible technique to alleviate I/R injury. The timing of LV unloading initiation is under investigation and the mechanisms associated with cardioprotection are largely unknown. Our data suggest that LV unloading simultaneously with coronary reperfusion leads to the least possible myocardial infarct size and most profound myocardial salvage. This improvement in myocardial salvage was associated with an upregulation of cardioprotective pathways including the expression of *PPARG*. Administration of rosiglitazone, a *PPARG* agonist, led to increased myocardial salvage following I/R injury. Future research could focus on interventions combining LV unloading with *PPARG* agonists to minimize I/R injury and maximize myocardial salvage.

### Limitations

We controlled for infarct size, but the ischemia duration that was used in our pre-clinical animal models, 30 minutes (mouse) and 90 minutes (swine) may not reflect the longer ischemic periods that patients might experience. Additionally, anatomical differences between swine and humans should be noted. Variable collateral circulation in swine might lead to inconsistent reperfusion outcomes, and thus the study of ischemia and reperfusion in pre-clinical animal models might not be equivalent to human pathophysiology.

## Supporting information

Supplement_Visker 2025

## AUTHOR CONTRIBUTIONS

JRV, ET, CPK, RH, MY, JL, TSS, EK, KS, EM, SN, AT, TTH, RMS, CL, FGW, and SGD were involved with the conceptualization of this manuscript. JRV, ET, CPK, RH, MY, JL, TSS, EK, LCR, JNV, KS, HK, YH, EY, EM, HS, LP, GP, SN, AT, GSD, JR, TTH, RMS, CL, FGW, and SGD were involved in methodology, validation, formal analysis, and invetigation. GSD, JR, TTH, RMS, CL, FGW, and SGD contributed reagents and resources. JRV, ET, CPK, RH, MY, JL, TSS, EK, LCR, JNV, KS, HK, YH, EY, EM, HS, LP, and GP were invovled in data curation. Writing the original draft of the manuscript was completed by JRV, ET, CPK, RH, LCR, CL, and SGD. Writing, reviewing, and editing was completed by JRV, ET, CPK, RH, MY, JL, TSS, EK, LCR, JNV, KS, HK, YH, EY, EM, HS, LP, GP, SN, AT, GSD, JR, TTH, RMS, CL, FGW, and SGD. CL, FGW, and SGD supervised the project.

## ACKNOWLEDGEMENTS

We would like to thank Chris Stubben for his analysis of the transcriptomics data. We would also thank the veterinarian staff at the University of Utah for their expertise in working with large animals. We would like to tahnk the University of Utah Metabolic Phenotyping Core, the Nora Eccles Harrison Cardiovascular Research and Training Institute, and the Nora Eccles Harrison Treadwell foundation for their support of this investigation. We acknowledge funding from the following sources: the National Institute of Health (NIH) under Ruth L. Kirschstein National Research Service Award T32HL007576 from the National Heart, Lung and Blood Institute, award number 3R35GM131854-04S1 to LCR, and award number 3R35GM1311854-04 to JR from the National Institute of General Medicine Sciences. We also would like to thanks the American Heart Association for Postdoctoral Fellowship Award 23Post1019351 to TSS and the Burrough Wellcome Fund for Postdoctoral Diversity Enrichment Program award to LCR. Next, we would like to thank the H.A. and Edna Benning Society at the University of Utah for funding contributed to SGD. JR is also an investigator of the Howard Hughes Medical Institute. The content of this manuscript is solely the responsibility of the authors and does not necessarily reflect the views of the NIH.

