## Supplement_Visker 2025 for "Peroxisome proliferator-activated receptor gamma (*PPARG*)-mediated myocardial salvage in acute myocardial infarction managed with left ventricular unloading and coronary reperfusion"

**SUPPLEMENTAL DATA**

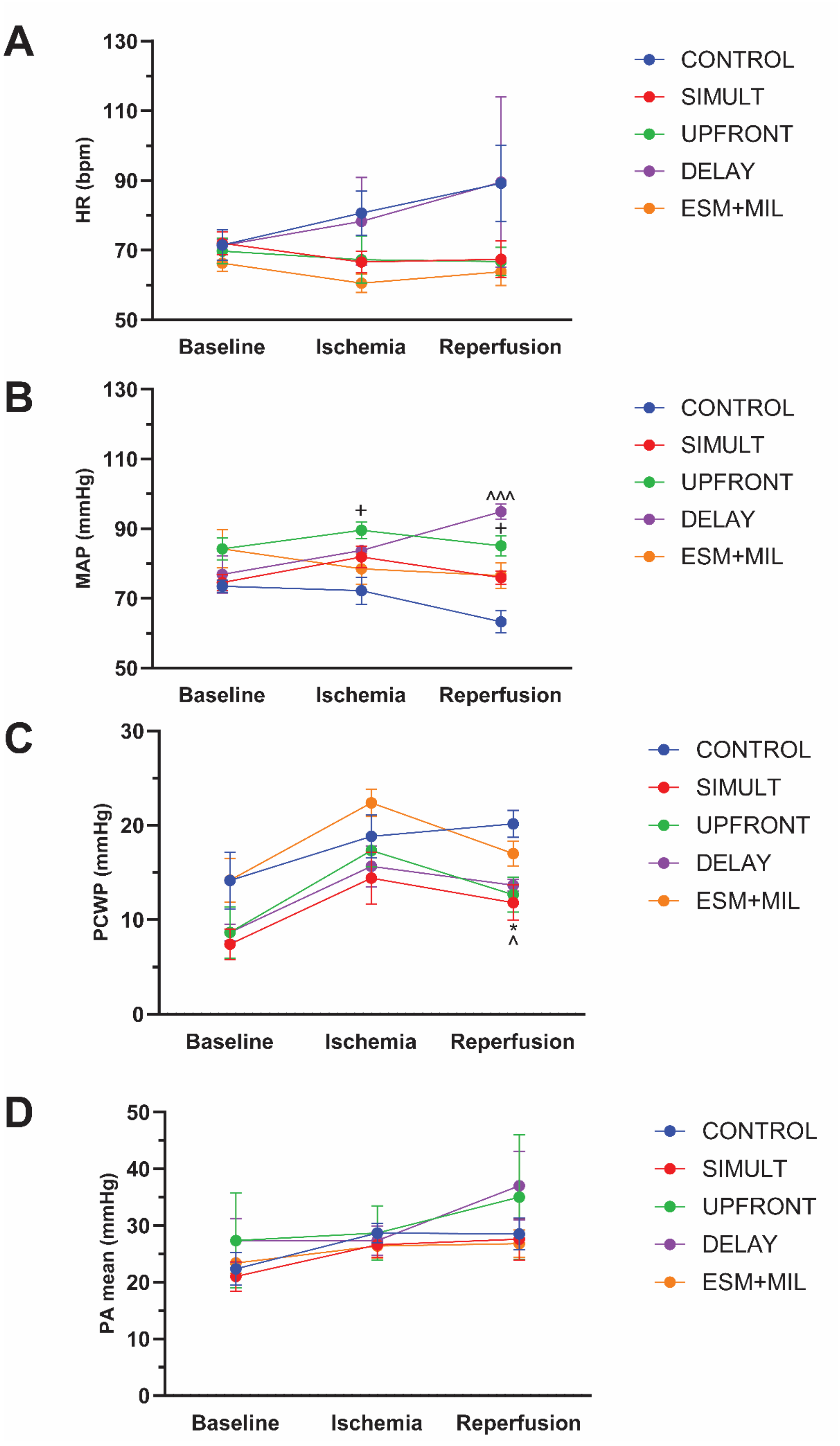

**Supplemental Figure 1. Cardiac performance measured by invasive hemodynamics during I/R injury.** LV-unloading sustained heart rate and arterial blood pressure within normal range, and reduced pulmonary capillary wedge pressure following acute I/R injury. **A)** Heart rate, **B)** Mean arterial pressure (MAP, mmHg), **C)** Pulmonary capillary wedge pressure (PCWP, mmHg), **D)** Mean pulmonary arterial pressure (PA, mmHg). A 2-way ANOVA with multiple comparisons (Tukeys HSD Post Hoc Test) was used compare the main effects of unloading (CONTROL, SIMULT, UPFRONT, DELAY, or ESM+MIL) and time (Baseline, Ischemia, and Reperfusion). + = $p<0.05$, CONTROL vs. UPFRONT. ^ = $p<0.05$, ^^^ = $p<0.001$, CONTROL vs. DELAY. * = $p<0.05$, CONTROL vs. SIMULT.
